# Alpaca Single B Cell Interrogation and Heavy-Chain-Only Antibody Discovery on an Optofluidic Platform

**DOI:** 10.1101/2023.02.10.528050

**Authors:** Mariya B Shapiro, Jacqueline Boucher, Anna Brousseau, Amin Dehkharghani, Justin Gabriel, Vishal Kamat, Ketan Patil, Feng Gao, Jennifer Walker, Ryan Kelly, Colby A Souders

## Abstract

*In vivo* discovery approaches for single domain antibodies such as VHH have been limited by the lack of methodologies available for camelid B cell interrogation. Here, we report a novel application of the Berkeley Lights Beacon® optofluidic platform to the discovery of heavy-chain-only antibodies by screening single B cells from alpacas immunized with two different targets. Custom methods for alpaca B cell enrichment, culture, on-Beacon IgG2/3 detection, and sequencing were developed and used to discover target-specific VHH candidates from an alpaca immunized with either human prostate specific membrane antigen (PSMA) or a second blinded target, in a proof-of-concept study. PSMA-specific VHH hits discovered on the Beacon were recombinantly expressed as VHH-Fc, purified, and characterized using label-free techniques. All but one VHH-Fc bound PSMA with a single-digit nanomolar affinity, and four candidates were successfully humanized *in silico* using a rapid bulk humanization approach. In addition, next-generation repertoire sequencing was performed at longitudinal timepoints after immunization, uncovering additional variants within the clonal lineages of the validated hits discovered on the Beacon platform. The establishment of this single B cell VHH discovery workflow extends the powerful Beacon technology to enable rapid discovery of VHH directly from natural camelid immune repertoires.

## INTRODUCTION

Antibodies typically consist of two heavy chains and two light chains, with both chains contributing to antigen binding. Over the past few decades, monoclonal antibodies have ushered in a new era of precision medicine. The field has evolved to encompass a broader range of biologic drug formats, opening the door for new modalities of immunotherapy. Due to their diverse structural properties, different biologic formats have distinct advantages and disadvantages with regards to biodistribution, *in vitro* and *in vivo* stability, and biological functions such as target specificity and engagement, which dictate their use in the treatment of disease.^1^

In addition to the standard antibody classes containing both heavy and light chains, camelids and some cartilaginous fish species also produce heavy-chain-only antibodies (HCAb) which are uniquely able to recognize their cognate antigen without a light chain.^2^ In camelids, the HCAb subclasses IgG2 and IgG3 consist of a variable domain (VHH) linked via a hinge to the Fc domain comprised of two constant regions (CH2 and CH3). Due to advantages such as their small size, high solubility and stability, relatively long CDRH3s, and ease of engineering, recombinant VHH domains—also known as single domain antibodies—have emerged over the past three decades^3^ as a promising biologic format for numerous applications in biomedicine.^4^ The drug caplacizumab^5^ became the first FDA-approved nanobody therapeutic in 2019, invigorating the VHH industry. Potential uses of VHHs have multiplied to include multispecific antibody constructs^6,7^ and chimeric antigen receptors (CARs)^8–11^, engagement of difficult targets^12,13^, biodistribution and localization studies^14–16^, and reagents for drug discovery^17^, with diverse therapeutic indications ranging from oncology^6–11,14–16^ to neurologic^18–20^ and infectious disease.^4,21,22^

Several platforms currently exist for single domain antibody (sdAb) discovery, each with distinct advantages. *In vitro* display technologies are widely used and involve the construction of recombinant libraries using VHH sequences from naïve or immunized camelids or sharks. These libraries are cloned into phage or yeast for surface display, and hits are discovered through iterative rounds of panning against a target antigen. While relatively cheap and rapid, these approaches have an increased risk for off-target reactivity and developability issues, are limited by the difficulty of using antigen formats other than recombinant proteins and depend heavily on the quality of the library and selection reagents. To overcome the obstacles associated with *in vitro* discovery, transgenic mice have been developed that express HCAbs because of CH1 exon deletion^23–25^. However, their usefulness is limited by the difficulty and long timeframe involved in generating these transgenic mice, as well as potentially weakened immune responses and reduced antibody affinity in some models^24^.

Natural *in vivo* VHH discovery in species expressing native HCAbs (e.g. alpacas), while technically more challenging and costly, has major advantages compared with *in vitro* display and transgenic murine models. Being larger animals, alpacas allow for large volume bleeds and repeated longitudinal sampling, resulting in a higher total potential screening throughput compared to smaller species like mice. Importantly, direct interrogation of HCAbs produced during the natural antibody response in alpacas may increase the chance of identifying antigen-specific candidates with higher affinity, lower off-target reactivity, and more desirable biophysical properties compared with alternative technologies currently available.

Compared with original approaches such as hybridoma and single B cell sorting, multiple next generation antibody discovery technologies have accelerated the discovery of therapeutic molecules. Single B cell screening platforms such as the Berkeley Lights Beacon combine relatively high throughput with complex screening strategies using multiplexed assays to enable antibody discovery against challenging targets ^26–29^. There is a growing need for next-generation antibody discovery methods that can be applied to VHH campaigns. However, the lack of validated phenotyping markers and reagents for enrichment and screening of desired camelid cell populations have posed obstacles for applying next generation technologies to *in vivo* VHH discovery. The Beacon has been used extensively for IgG antibody discovery using mouse and human samples. However, to our knowledge, the advantages of this groundbreaking technology have not yet been extended to camelids. In this study, we establish methods for enriching and culturing primary HCAb-secreting B cells from immunized alpacas and use the Beacon optofluidic platform to screen and sequence antigen-specific HCAbs. We present the results of our proof-of-concept case study using prostate-specific membrane antigen (PSMA) as a model target for *in vivo* VHH discovery and compare single cell screening results to a target antigen that did not elicit a detectable HCAb serum response. A subset of VHH sequences were chosen for recombinant expression, and the resulting VHH-Fc constructs were characterized by measuring their PSMA binding affinity by surface plasmon resonance (SPR) while their epitope diversity was assessed using biolayer interferometry (BLI). Clonotype lineages related to the characterized VHHs were longitudinally analyzed by repertoire sequencing throughout the 216-day immunization process. Finally, the VHHs were humanized using a rapid and high throughput *in silico* method.

## METHODS

### Alpaca immunizations and tissue harvests

One alpaca was immunized with PSMA-His protein (Sino Biological) and CFA/IFA adjuvant administered every 21 days over the study period. PBMC harvests for Beacon screens were timed to occur 10 days after the most recent boost. Two additional alpacas were immunized with Target B: one alpaca received a recombinant protein-based regimen analogous to the PSMA immunization protocol, and the other received a primarily-DNA-based immunization strategy up to day 178, and received protein boosts thereafter. Beginning on day 73 of immunization for the anti-PSMA alpaca and day 94 for the anti-Target B alpacas, peripheral blood samples were collected 10 days after the previous boost, at intervals spanning up to 9 months of sampling. PBMC isolated from 300mL of peripheral blood were used as input for the magnetic enrichment of memory B cells.

### Serum titer tests

For the PSMA study, titer tests were performed via indirect ELISA to human PSMA-His and murine PSMA-His (Sino). For the Target B study, titer tests were performed via indirect ELISA and Streptavidin Capture ELISA to AVI-HIS-Target B and Target B-AVI-HIS (generated in-house) and Human and Mouse HIS tagged Target B proteins (Sino). Anti-alpaca IgG2/3 or anti-alpaca IgG(H+L) secondaries were used to detect HCAb or total IgG in the serum respectively.

### Cell sample preparation and Beacon screening

PBMCs were isolated from whole blood using density-gradient centrifugation. Biotinylated enrichment reagents bound to streptavidin-coated paramagnetic beads were used for magnetic separation of memory B cells from the total PBMC population. These memory B cells were subsequently cultured over a period of several days in stimulation media to induce IgG secretion. On the day of screening, stimulated memory B cells were resuspended, counted and brought to the appropriate density for import on the Berkeley Lights Beacon OptoSelect Chip. Once imported, single live B cells were allocated into individual pens on the OptoSelect Chip using optoelectrical positioning (OEP). After individual B cells were penned, IgG capture beads and fluorophore-labeled IgG2/3 detection reagents were imported onto the chip and the chip was imaged over the course of 45 minutes. This assay was then flushed from the chip using B cell culture media. IgG capture beads and fluorophore-labeled recombinant PSMA were then imported onto the OptoSelect chip and imaged over the course of 45 minutes. For some experiments, assays were run concurrently with both PSMA and IgG2/3 detection reagents to simultaneously obtain both readouts. After assays were conducted, a combination of artificial intelligence-based and manual scoring methods were used to discern binary blooms (signal above background increasing over time).

### Sanger VHH sequencing

Cells of interest identified during the Beacon screen were exported into 96-well PCR plates containing lysis buffer and mineral oil. Heavy chain variable regions were amplified using gene specific primers for cDNA synthesis and two-step nested PCR. The forward and reverse primers bind to the leader and CH2 region of the heavy chain, respectively, and amplify IgG1, IgG2, and IgG3 isotypes. Amplicons were sent for Sanger Sequencing and data was analyzed in Geneious Biologics using a customized *Vicugna pacos* Single Clone Analysis Pipeline to determine antibody annotations. VHH (IgG2 and IgG3) antibodies were identified by signature sites in FR2 and the constant region. Full H+L (IgG1) antibodies were filtered out. VHH candidates of interest were chosen for recombinant expression, codon optimized, and cloned into an expression vector on a human IgG1 or murine IgG2a Fc backbone (Azenta).

### Hit Expansion (repertoire sequencing)

RNA was isolated from memory B cells (5e6-10e6 cells) enriched from PBMC at various timepoints. Sequences were amplified using gene specific primers for cDNA synthesis and PCR. RNA input was normalized and multiple thermocycling conditions were used to determine conditions for optimal amplification. Amplicons were sequenced by Next Generation Sequencing (Illumina). Sequences were annotated in PipeBio, using the *Vicugna pacos* germline. NGS reads with CDR combinations occurring only once in the NGS dataset were removed. NGS sequences similar to the single B cell clone sequences were obtained by co-clustering the NGS reads with the single B cell clone sequences using a CDR3 identity percentage of 80%. 19 of these sequences were removed due to the presence of stop codons. All unique VDJ sequences were subsequently extracted from the data, resulting in the 386 unique sequences shown in Figure 6.

### Recombinant expression of VHH Fc constructs

Expi293 HEK cells were transfected with plasmid containing either VHH-mouse IgG2a Fc (VHH-mFc) or VHH-human IgG1 Fc (VHH-hFc) expression cassettes via typical polyethylenimine methods. Transfected cells were treated with valproic acid and tryptone 24 hours after transfection. After 5 days in culture, supernatant was isolated from transfected cells and antibodies were purified from the supernatant using Protein A-based affinity purification.

### SPR binding affinity studies

VHH antibodies were assessed on a Carterra® LSA™ platform. All the binding studies were performed in running buffer containing 10mM HEPES, 150mM NaCl, 0.05% Tween20, pH7.4 at 25°C. Purified antibodies were first captured in an array format on the anti-human or anti-mouse Fc antibody surface. After 7-8 buffer injections, increasing concentrations of human PSMA (Sino) ranging from 2nM to 200nM were tested with a 10-minute association phase and 30-minute dissociation phase. The real-time binding data was analyzed using a double reference subtraction procedure on the Carterra kinetics software (version 1.8).

### BLI epitope binning studies

An epitope competition assay was performed via BLI on an Octet RH96 platform (Sartorius) using an in-tandem assay format. The entire experiment was performed at 25°C in 10mM HEPES, 150mM NaCl, 0.05% v/v Tween-20, 1mg/mL BSA, 0.02% NaN3, pH7.4 (HBS-BNT) buffer with the plate shaking at a speed of 1000rpm. To assess whether two antibodies compete with one another for binding to their respective epitopes, huPSMA-His was first captured onto anti-His antibody (HIS1K) coated Octet biosensor tips. The huPSMA-His captured biosensor tips were then saturated with the first PSMA antibody (subsequently referred to as mAb-1) by dipping into wells containing 50µg/mL solution of mAb-1 for 4 minutes. The biosensor tips were then subsequently dipped into wells containing 50µg/mL solution of second PSMA antibody (subsequently referred to as mAb-2) for 3 minutes. The biosensor tips were washed in HBS-EBT buffer in between every step of the experiment. At the end of each cycle the biosensors were regenerated in 10mM Gly, pH2.0 containing 150mM NaCl and the entire cycle was repeated for subsequent antibodies. Epitope binning studies were performed in both directions to assess competition profile based on the order of PSMA antibody binding. Epitope binning data was analyzed using the Octet data analysis HT (version 11.0) and Carterra epitope software (version 1.8) to generate clustered heat map and community network diagrams.

### High throughput in silico humanization

A machine learning-based pipeline was used for antibody design to enable bulk humanization. The platform recognizes non-human residues using deep learning attention mechanisms trained on human antibody sequences. The final humanized sequences were produced by taking the most probable predicted residue at each position, except in CDRs, where mutations were ignored and the original parental sequence was preserved. Five different designs based on fixed residue locations were used to generate one variant per definition for each parental VHH. Additional back-mutations in the framework region 2 were performed manually for stability of VHHs. Humanness percentage of parental and humanized sequences was calculated in Geneious Biologics (Human Ig May 16 2018 database) by pairwise comparison of VDJ regions against the closest human germline sequence.

## RESULTS

A uniquely powerful tool for antibody discovery, the Berkeley Lights Beacon enables screening of antibodies secreted from single B cells for biologically relevant functions such as target binding, ligand blocking, and inhibition of signal transduction. The Beacon technology uses structured light to segregate antibody-secreting cells into thousands of individual nanopens on an OptoSelect chip in a process called ‘optoelectrical positioning’. Each nanopen holds a single B cell and opens at one end into a common channel. As the secreted antibodies continuously diffuse from the nanopen into the channel, their functional antigen-binding properties are measured using time-lapse fluorescent imaging assays containing capture beads, fluorescently-labeled target molecules, target-expressing cells, or other assays. A spreading “bloom” of fluorescent signal in the channel area above the source pen is typically considered as a positive result. After the completion of each assay, the channel can be flushed out and a new assay is imported. Multiple assays can be performed sequentially on a single B cell and different functional attributes of the antibodies can be assessed. After the assay results are analyzed, the B cells of interest can be individually exported and the mRNA encoding the IgG can be sequenced.

By utilizing screening strategies involving diverse reagent suites and binding orientations, our lab has established new assay formats to provide valuable functional data at the single cell stage of the antibody screening process, enabling efficient down-selection of functional hits for recombinant expression and validation. Several examples of custom functional assays are illustrated in Figure 1. In the first example, a custom functional assay was developed to identify antibodies capable of blocking a receptor-ligand interaction that leads to an intracellular signaling cascade (Figure 1A). This assay design relied upon a cell line in which, upon ligand binding to a receptor, triggers a signaling cascade that leads to the secretion of a reporter molecule. The reporter cells were co-penned and incubated together with antibody-secreting B cells prior to ligand exposure. An assay was subsequently imported in the channel to capture and detect the secreted reporter molecule. Active signaling through the receptor was indicated by the fluorescent detection of secreted reporter in the channel above the source pen. In contrast, blocked signaling was distinguished by the absence of a fluorescent signal in the channel above the source pen, indicating a lack of reporter secretion. The second example shows a pair of cell surface binding assays, in which secreted antibodies bind to target cells in the channel and are detected using a fluorescently labeled secondary reagent (Figure 1B). By carefully choosing the target-positive and counter-screen cell lines, this screening strategy can be used to identify target-specific cell binders. The final example illustrates a ligand blocking assay performed using fluorescently labeled recombinant proteins (Figure 1C). In this setup, the ligand and target are each labeled with a different fluorophore and imported into the channel with IgG capture beads. A secreted antibody that blocks the interaction will be identified by positive signal from the fluorophore corresponding to the target only, while a non-blocker will appear as a positive signal from both fluorophores indicating simultaneous binding of the antibody and the ligand to the target protein. The assay formats described above can be performed sequentially during a single Beacon screening day, providing an unparalleled level of detail regarding antibody function for more than ten thousand single B cells per Beacon chip.

**Figure 1.**
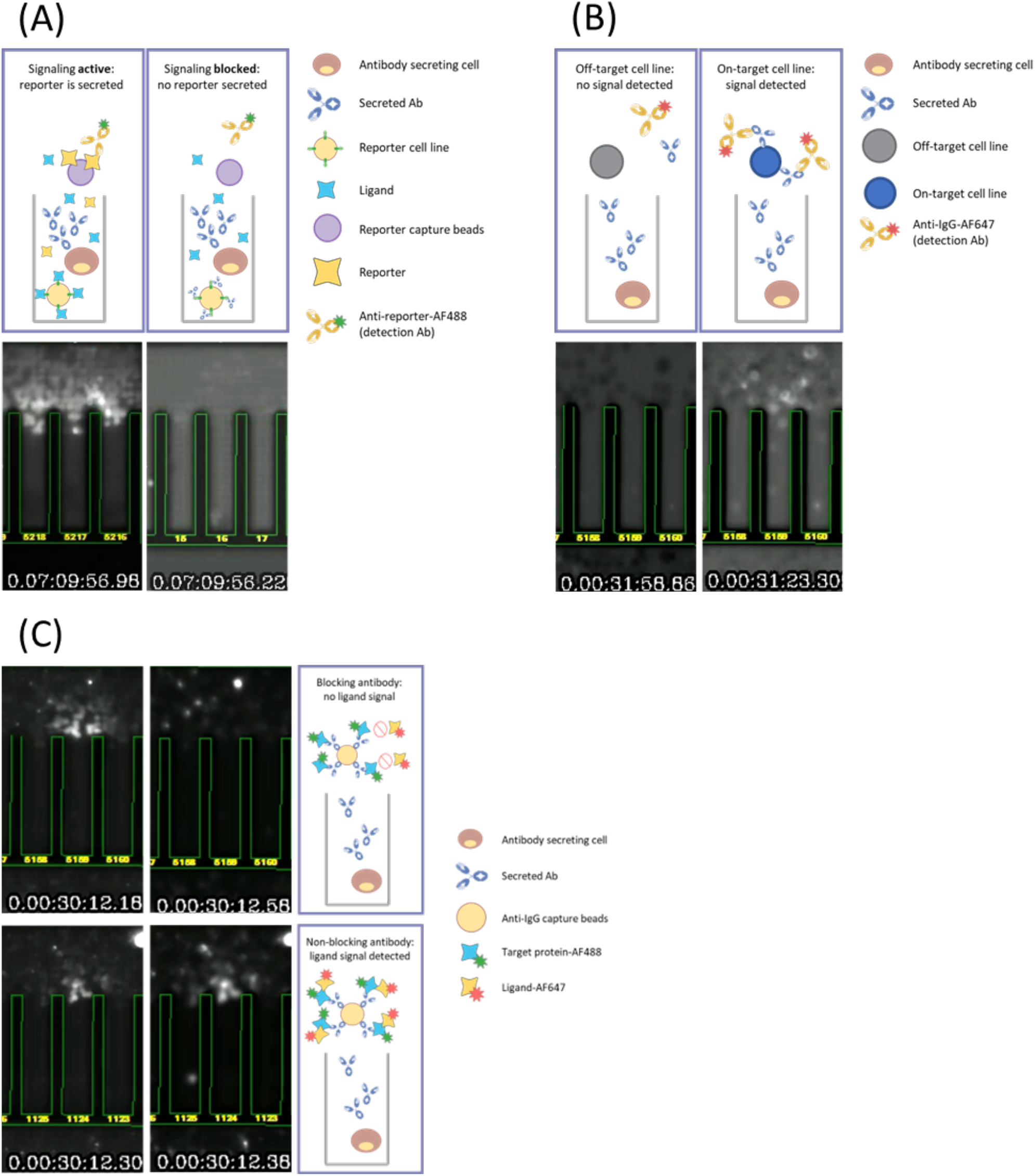
Functional assays for Beacon screening to determine biologically relevant properties of antibodies secreted from individual B cells. (A) Cell-based reporter assay. Reporter cells are sequestered with B cells inside nanopens. Ligand-receptor binding drives signaling leading to secretion of a reporter molecule that is captured on beads and detected by a fluorophore-labeled antibody. (B) Cell binding assays. Target cells and fluorophore-labeled detection antibody are imported to interrogate antibodies secreted by B cells. Specific hits show binding to on-target cells but not off-target cells in sequential assays. (C) Ligand blocking assay. Target and ligand, labeled with different fluorophores, are imported with anti-IgG capture beads. Antibodies secreted by B cells are captured on the beads and encounter target-ligand complexes. A blocking antibody outcompetes the ligand for binding to the target, resulting in fluorescent signal from the target only. A non-blocker binds the complex, resulting in signal from both reagents.

To address whether the Beacon platform could be adapted for screening primary B cells from camelids, we decided to use alpaca (*Vicugna pacos*) as a model for *in vivo* HCAb discovery. In this study, immunized alpacas were used as a source of antibody-secreting B cells for development and optimization of the Beacon screening workflow (Figure 2). We used prostate-specific membrane antigen (PSMA) as a model target antigen for this study. In addition, we investigated screening of alpaca B cells following immunization regimens against Target B, another cell surface target. However, unlike PSMA, Target B did not elicit a detectable IgG2/3 isotype response, and therefore antigen-specific IgG2/3 B cells were expected to represent very rare events.

**Figure 2.**
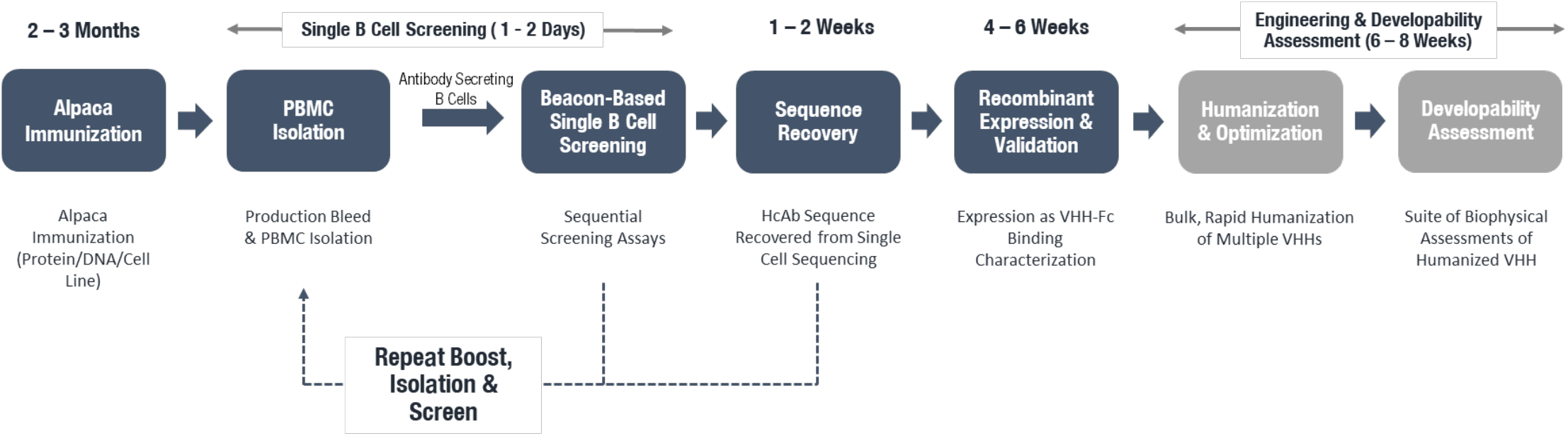
Overview of the alpaca single B cell screening and VHH discovery workflow. Alpacas are immunized for a minimum of 2-3 months and up to 9 months before proceeding to PBMC isolation and subsequent single B cell screening. Multiple rounds of screening can be performed following additional immunization boosts. Downstream recombinant expression, humanization, and validation provide data to enable lead candidate selection. Immunization, screening, and sequencing were performed for both the PSMA and Target B campaigns, while recombinant expression, humanization, and validation were only performed for the PSMA campaign. Developability assessment was not performed for the campaigns in this study.

The goal of alpaca immunization was to achieve a strong HCAb serum titer against the targets. To understand the kinetics of the antibody response in alpacas, total antigen-specific IgG was measured, along with target-specific and total (target agnostic) IgG2/3 titers at different timepoints by ELISA (Figure 3). In the alpaca immunized with PSMA, the total IgG titer against the target reached an EC_50_ of approximately 1:5400 serum dilution and remained unchanged from day 97 through day 174. In contrast, IgG2/3 titer gradually increased throughout the study and reached a maximum EC_50_ of approximately 1:900 by day 153 (Figure 3A). Interestingly, a distinct antibody response was observed in a second and third alpaca which were immunized with Target B recombinant protein or DNA, with total IgG against the target gradually increasing over time; however, no IgG2/3 titer was detectable at any point in the study in either alpaca (Figure 3B). The difference between alpacas immunized with distinct antigens suggests that the target identity and potential variation between individual animals may contribute to differences in the kinetics of the antibody response. Notably, total IgG2/3 titers (antigen-agnostic) in the three animals were broadly similar and changed very little over the course of the study (Figure 3C), indicating that a difference in antigen-specific IgG2/3 titer is not attributable to a deficient total IgG2/3 response across alpacas.

**Figure 3.**
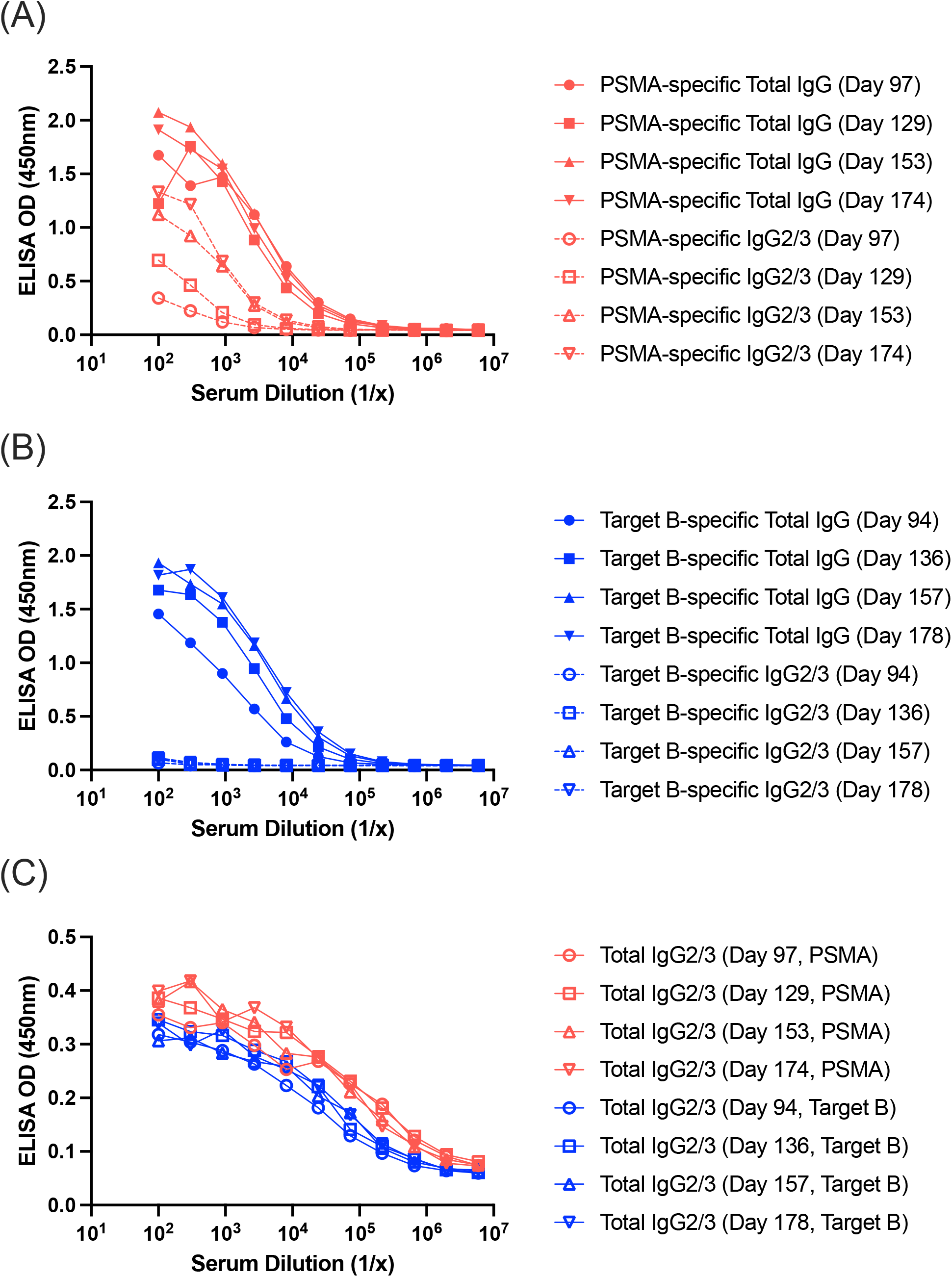
Serum titer following alpaca immunization with PSMA or Target B. (A) PSMA-specific titers in alpaca serum. (B) Target B-specific titers in alpaca serum. Total target-specific IgG titer is depicted by solid lines while HCAb (IgG2/3 isotype) target-specific titer is depicted by dashed lines. (C) Total HCAb (IgG2/3 isotype) titer independent of antigen specificity is compared across all immunized alpacas. Each data point shown for Target B is an average for the two alpacas immunized with this target.

To enable Beacon-based identification of desired target-specific candidates, novel methods were developed for enriching, culturing, and detecting HCAb-secreting B cells that bind the target (Figure 4). On the day of each alpaca production bleed, PBMC were processed to enrich alpaca memory B cells, which were then stimulated in culture using an optimized method to stimulate IgG secretion. The methods involved in these procedures were developed over the course of >50 screening experiments designed to test variables such as the concentration and type of enrichment reagent, composition of the culture media, duration of incubation in culture, and sample processing conditions prior to import onto the Beacon. On the day of the Beacon screen, the cultured B cells were imported onto the Beacon chip, and single cells were segregated into nanopens in preparation for analysis. Custom anti-alpaca IgG capture beads were generated and, after premixing with anti-alpaca IgG2/3-AF488 secondary antibody, imported into the channels of the Beacon chip. Longitudinal imaging in the FITC channel visualized the signal from the capture and detection of secreted alpaca IgG2 and IgG3 subclasses (Figure 4A). Following this first assay, chips were flushed and a second assay performed in which the anti-alpaca secondary was replaced with AF647-labeled recombinant PSMA protein. In this format, the secreted antibodies were captured on beads and binding to the fluorescent PSMA antigen produced a positive signal above the pen (Figure 4B). Desired hits were chosen for export and sequencing based on a positive readout in both assays. The same approach was applied for Beacon screens on Target B, replacing the fluorescently labeled PSMA in Assay 2 with the antigen of interest for that campaign.

**Figure 4.**
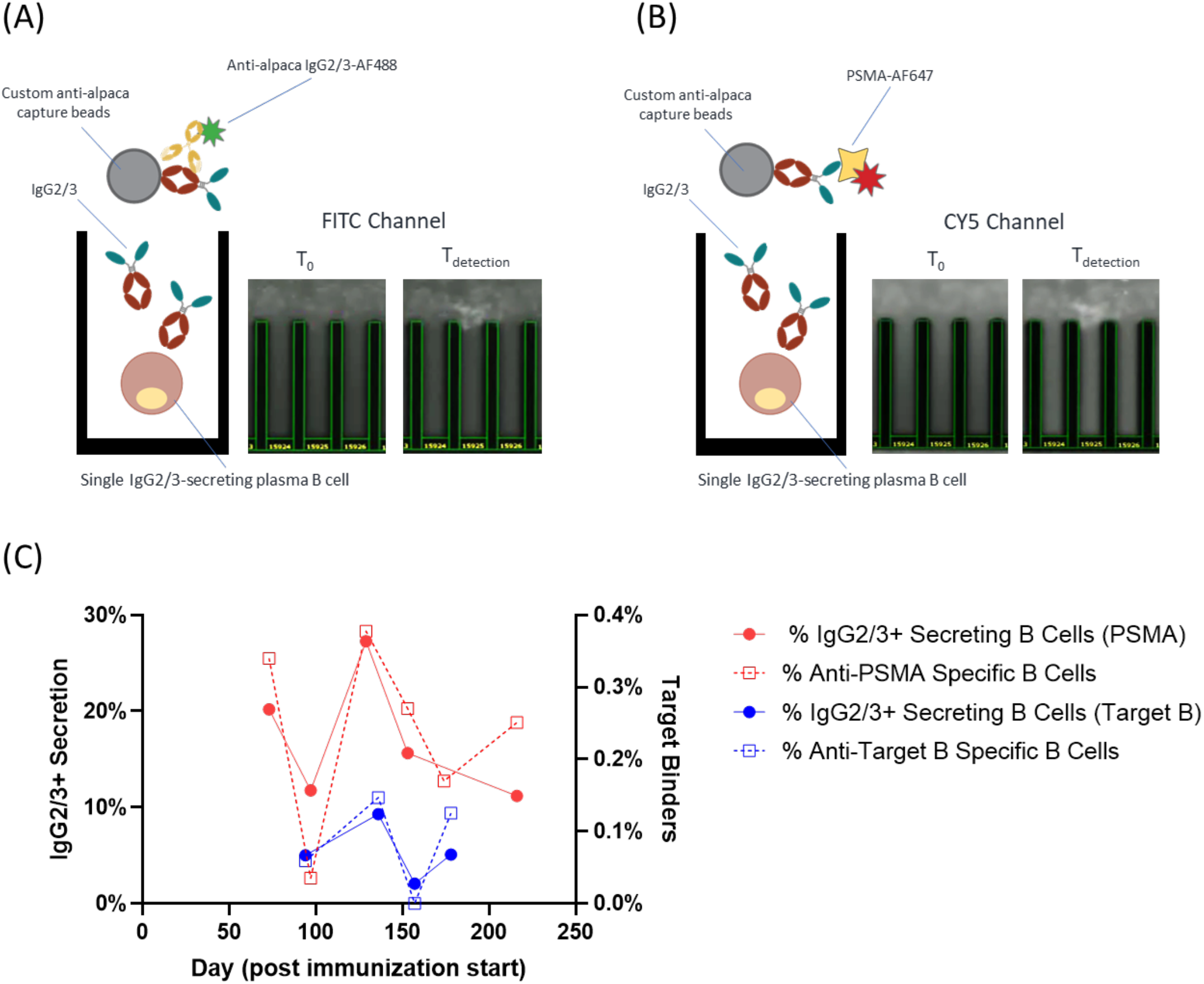
Beacon screening strategy and longitudinal assay results. (A) Assay to detect secretion of IgG2 and IgG3 antibodies. Alpaca IgG capture beads are imported alongside an AF488-labeled antibody specific for IgG2/3 to interrogate secreted antibodies, followed by imaging in the FITC channel on Beacon. (B) Assay to detect PSMA-specific IgG secretion. Alpaca IgG capture beads are imported alongside AF647-labeled PSMA, followed by imaging in the CY5 channel on Beacon. Similar assays were used for Target B. Assays were either performed sequentially, or concurrently with readouts in both channels indicating IgG2/3 secretion and target binding respectively. Representative images show a positive hit. (C) IgG2/3 secretion assay results (solid lines, left y-axis) and target-specific assay results (dashed lines, right y-axis) were measured on Beacon at indicated time points following initial immunization. To account for screens involving pooled alpaca B cell samples, each data point represents the average of all samples screened.

A distinct advantage of alpacas is their large size, which enables repeated longitudinal sampling and screening. To investigate the IgG2/3 secretion rates and target binding rates over time, Beacon screening workflows were repeated using PBMCs collected at different time points for each PSMA and Target B immunized alpaca. IgG2/3 secretion rates for the PSMA immunized alpaca, which elicited a robust anti-PSMA IgG2/3 titer, followed a consistent pattern with HCAb secretion rates reaching a peak of ∼30% around 130-140 days after the start of immunization and declining somewhat thereafter, but remaining above 10% even beyond 200 days (Figure 4C). As expected, target binding rates were considerably lower than overall IgG2/3 secretion rates; however, the longitudinal pattern tracked closely with the total HCAb secretion rates. The PSMA-immunized alpaca had higher IgG2/3 secretion and target binding rates as compared to the Target B-immunized alpacas, in agreement with the IgG2/3 serum titers. Interestingly, IgG2/3 secreting and target-specific B cells were still observed in Target B-immunized alpacas despite the lack of detectable serum titer, and 51 unique VHH sequences were recovered during this campaign.

To characterize the antibodies discovered during the PSMA Beacon screens, a subset of 13 B cell clones were selected and recombinantly expressed as VHH-mFc. Based on the SPR binding studies, 12 out of 13 VHH-mFc bound PSMA with a single digit nanomolar affinity (Figure 5A, 5B; Supplementary Table 1). The cross-competition assay performed on BLI revealed that the 13 VHH-mFc recognized largely non-overlapping epitopes and eight epitope bins were identified (Figure 5C). Five epitope bins (bin# 2, 3, 6, 7, 8) were represented by just one VHH-mFc while three additional bins (bin# 1, 4, 5) were comprised of two or more VHH-mFc (Figure 5D).

**Figure 5.**
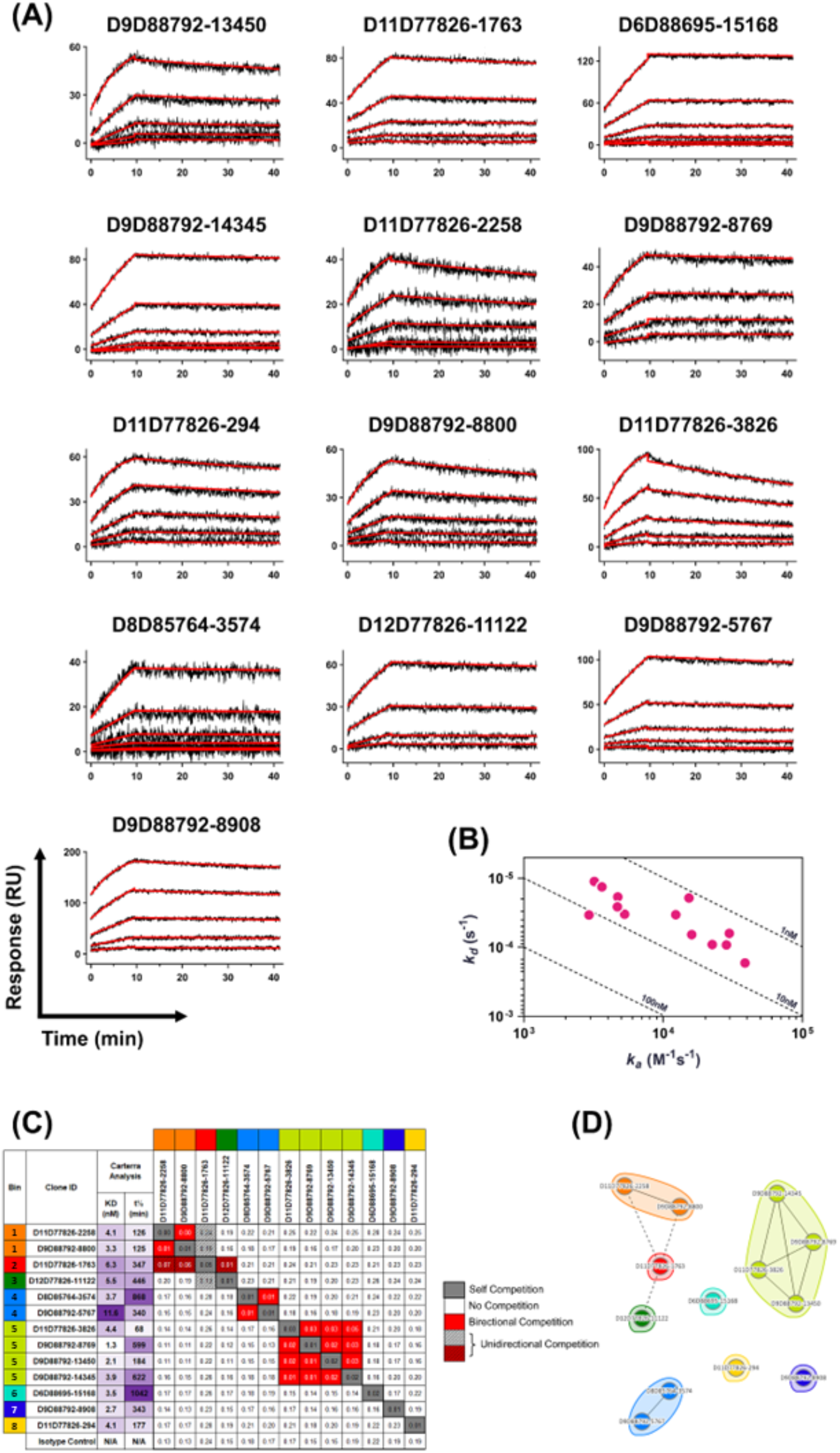
Characterization of a subset of VHH candidates identified by single B cell screening against PSMA. (A) Individual surface plasmon resonance sensorgrams of the 13 recombinantly expressed VHH-mFc antibodies binding to human PSMA-His. VHH-mFc were immobilized and sequentially increasing concentrations of PSMA-His in solution were flowed over the chip. (B) Isoaffinity plot summarizing the affinity data in part (A). (C) Characterization of binding epitopes by competition assay using biolayer interferometry. Eight distinct epitope bins were targeted by the 13 candidates. (D) Network map of epitope bins identified in the competition assay.

Although the Beacon offers a relatively high-throughput screening approach (∼10^4^ B cells per Beacon screen), these B cells represent a small fraction of the antibody repertoire of an immunized alpaca. In addition to sequencing the exported Beacon hits, sequences from the PSMA-immunized alpaca total B cell population were assessed on unused memory B cells enriched from PBMCs at various timepoints by analyzing the antibody repertoire via next generation sequencing (NGS). Using this approach, distinct clusters of sequences were identified that were closely related to validated antibodies discovered on the Beacon (Figure 6A). Depending on the cluster, closely related sequences were observed across two, three, or all four timepoints, suggesting a gradual evolution of the immune response in this alpaca with dominant clonal lineages expanding and contracting during the immunization regimen. Repertoire sequences related to single cell clones were present in 7, 9, 9 and 3 distinct clonotype lineages at days 73, 129, 153 and 216 post-immunization, respectively. Within the top three lineages of the repertoire sequences, a wide range of sequence liability scores and V gene identities was observed, providing a source of additional sequences of interest with potentially improved properties compared with their Beacon-discovered family members (Figure 6B). Sequence alignment of these closely related Beacon and repertoire clones revealed a high degree of diversity of individual point mutations in framework as well as CDR1 and CDR2 regions, suggesting possible point mutations for improving clone properties (Figure 6C).

**Figure 6.**
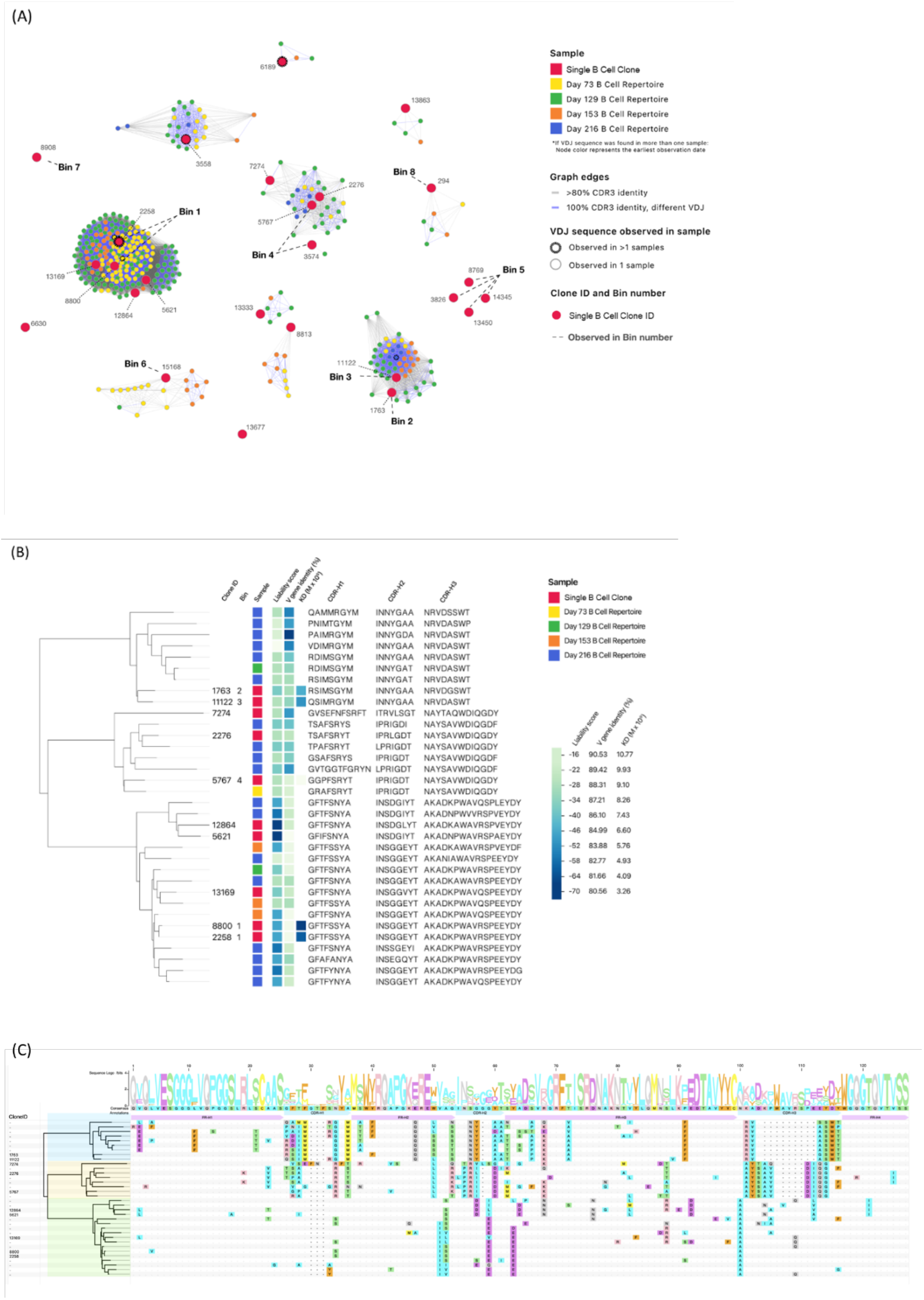
Relationship of total HCAb B cell repertoire sequences to VHH candidates isolated from single B cell screening from a PSMA-immunized alpaca. (A) Repertoire sequences from four different screening timepoints post initial immunization were grouped in relation to PSMA-specific single B cell candidates by >80% CDR3 identity (gray connecting line) or 100% CDR3 identity (blue connecting line). (B) Epitope bin is included as a characterization parameter to easily visualize relevant epitope relationships among clonal lineages. The top three clonal lineages by clonotype count were further analyzed to compare sequence liability score, V-gene identity, affinity (KD) and CDR amino acid sequences. (C) The same lineages were aligned across the entire VDJ segment to compare point mutations between single B cell candidates (Clone ID shown at left) and related repertoire sequences.

In addition, the parental VHH antibodies were humanized *in silico* (Table 1). The humanness of the starting parental clones ranged from 68.0 to 82.7% (average 73.9%). After humanization, average humanness per candidate increased by 9.2% ± 3.3% (range = 1.9% to 16.3%), while the affinity of humanized candidates decreased by a median of 2.35-fold as compared to parental affinity, depending on the humanization method utilized. At the extremes of the five humanization designs tested, design 1 was the least aggressive with respect to humanness score (average humanness improvement of 6.2% ± 2.0%), but it also preserved the highest fidelity binding affinity (median decrease in K_D_ of humanized candidates was 1.85-fold). Conversely, design 5 was the most aggressive in terms of achieving the highest % humanness (average humanness improvement of 11.5% ± 3.3%), but also resulted in 90% of clones losing all binding to the target after humanization. Importantly, many of the humanized variants achieved substantial increases in humanness with only modest reductions in target binding affinity. For example, humanization of candidate D11D77826-2258 using design 3 resulted in a 10.2% increase in humanness (from 82.7% to 92.9%) with only a 2.1-fold change in target binding affinity. Ultimately, a gradation of humanization aggressiveness revealed no single standard was universally superior. The optimal method to balance humanness with preserved binding characteristics varied across candidates, underscoring the value of testing different humanization designs followed by empirically measuring target binding affinity for each resulting variant to generate the optimal humanized VHH lead candidate. Using acceptance criteria of 80% or greater humanness and less than three-fold change in affinity, four of 12 candidates were successfully humanized.

**Table 1.** Humanized VHH candidate humanness percentage and fold-change in binding affinity across five humanization designs, as compared to parental sequences. *N.T. = Not Tested; ^#^N.B. = No Binding; ^$^I.C. = Binding detected, but inconclusive affinity due to poor curve fit analysis.

| Clone # | Parental | Method 1 |  | Method 2 |  | Method 3 |  | Method 4 |  | Method 5 |  |
| --- | --- | --- | --- | --- | --- | --- | --- | --- | --- | --- | --- |
|  | % Humaness | % Humaness | Fold change in KD | % Humaness | Fold change in KD | % Humaness | Fold change in KD | % Humaness | Fold change in KD | % Humaness | Fold change in KD |
| D11D77826-1763 | 73.5 | 80.6 | 1.0 | 81.6 | 2.3 | 82.5 | 3.9 | 84.7 | N.B. # | 85.6 | N.B. # |
| D11D77826-2258 | 82.7 | 88.8 | 1.7 | 92.9 | 2.1 | 93.9 | N.T. * | 90.8 | 1.7 | 93.9 | N.B. # |
| D11D77826-294 | 74.5 | 81.6 | 9.4 | 84.7 | 2988 | 85.7 | N.B. # | 84.7 | N.B. # | 86.7 | N.B. # |
| D11D77826-3826 | 69.4 | 77.6 | 2.4 | 81.6 | 73 | 82.7 | 143 | 84.5 | 27 | 85.7 | N.B. # |
| D12D77826-11122 | 71.4 | 79.6 | 6.7 | 80.6 | 1.3 | 81.4 | 8.7 | 83.7 | N.B. # | 84.5 | N.B. # |
| D6D88695-15168 | 74 | 77.0 | 1.3 | 77.0 | 1.9 | 77.0 | N.T. * | 80.0 | N.B. # | 78.0 | 1.0 |
| D8D85764-3574 | 73.5 | 77.6 | 1.6 | 81.6 | 18 | 83.7 | N.B. # | 82.7 | I.C. \$ | 83.7 | N.B. # |
| D9D88792-13450 | 74.5 | 81.6 | 11 | 83.7 | 5.1 | 84.7 | 22 | 85.6 | N.B. # | 85.6 | N.B. # |
| D9D88792-5767 | 74.5 | 81.6 | 1.2 | 82.7 | 1.3 | 84.5 | 1.7 | 86.7 | N.B. # | 86.7 | N.B. # |
| D9D88792-8769 | 68.0 | 75.3 | 3.5 | 77.3 | 17 | 79.4 | N.B. # | 79.4 | I.C. \$ | 83.5 | N.B. # |
| D9D88792-8800 | 80.6 | 87.8 | 1.2 | 92.9 | N.B. # | 93.9 | N.T. * | 90.8 | N.T. * | 93.9 | N.T. * |
| D9D88792-8908 | 70.3 | 72.2 | 2.0 | 74.2 | N.B. # | 73.2 | N.B. # | 75.2 | N.B. # | 76.7 | N.B. # |

## DISCUSSION

Adoption of high-resolution screening assays during the early antibody discovery workflow allows rapid identification of valuable hits and can accelerate therapeutic antibody discovery. However, such approaches have not been widely implemented for single domain (e.g. VHH) antibody discovery. For the first time, in this study, we established a novel platform for function-forward VHH discovery by pairing high-resolution Beacon screening capabilities— which have been successfully used for immunized mice^27,29^ and human donors^26^—with a newly developed upstream workflow for alpaca B cell enrichment, stimulation, and screening. This platform was used to discover and validate a panel of 13 antibodies that bound PSMA with nanomolar affinity and exhibited a broad epitopic coverage and sequence diversity. Moreover, repertoire sequencing identified additional mutations and closely related variants. Finally, a rapid *in silico* high throughput humanization approach resulted in multiple lead candidates with humanness scores at acceptable levels for clinical studies and preserved binding affinity. Indeed, the success of different humanization designs for different candidates supports the value of library-based approaches to humanization paired with affinity maturation for optimizing top candidates. Taken together, to our knowledge we propose the most rapid high-resolution workflow for discovery and engineering of VHH candidates from a natural immune response.

The Beacon and other nanowell-based microfluidic instruments operate by enabling the interrogation of antibodies secreted by antibody-secreting cells into the channel. Therefore, when starting with a PBMC population, optimizing isolation of B cells and stimulation culture conditions is essential for achieving a sufficient IgG secretion rate to support effective Beacon screening. With on-Beacon IgG2/3 secretion rates ranging from 3-30% depending on the alpaca, immunogen, serum response and timepoint, our rates are comparable to IgG secretion rates typically seen in our immunized murine plasma cell screens (interquartile range 2.1-22%, unpublished data), supporting that the alpaca B cell isolation and activation process performs similarly to standard processes in well-established species. Encouragingly, the methods established here can lead to successful antigen-specific HCAb identification even from alpacas that appear to have no detectible IgG2/3 antigen-specific serum titer response, as demonstrated in our campaign against Target B.

Analysis of results from single B cell screening and longitudinal repertoire sequencing suggests ideal diversity and quantity of antigen-specific candidates appears approximately 4-5 months from the start of immunization. Consistent across two different antigens, the peak IgG2/3 and antigen-specific single B cell response was observed at 129 and 136 days post-immunization. In conjunction, repertoire sequences from the individual case example represented the highest proportion of single B cell clone lineages at the same timeframe, suggesting that antigen-specific clonal expansion is possibly maximized at this time. The combination of single cell validation data with repertoire data provides a unique perspective to filtering repertoire data to relevant lineages more likely to be antigen-specific, as opposed to full repertoire sequencing alone that predominantly reports irrelevant binding clones. While the data is only from a single alpaca and requires validation with additional studies, we propose that the identification of clonal diversity in the present study is more accurate as compared to solely repertoire sequencing and has potential utility in generating antigen-biased immune libraries for additional screening. Our findings may inform future studies to select optimal immunization timepoints for PBMC harvest to produce immune libraries and/or single B cell screening.

In this study, we primarily focused on interrogating the memory B cell compartment in alpacas by activating these cells in culture. Future research should expand the screening capabilities to other relevant B cell subsets, especially plasma cells. However, a lack of validated enrichment reagents and cell population markers pose challenges for such efforts, and additional basic research to identify relevant cell surface biomarkers of alpaca B cells would provide a strong foundation for these innovations.

The goal of the present study was to develop and optimize methods for alpaca B cell enrichment, activation, and HCAb detection on the Beacon. A major limitation of this study is the small number of alpacas used in the experiments described here: one alpaca immunized with PSMA and two alpacas with Target B. Of the two alpacas immunized with Target B, one was immunized with a similar recombinant protein-based regimen as the PSMA alpaca, while the other was immunized using a distinct DNA-based regimen. Therefore, these three alpacas cannot be compared directly, and the IgG titer and secretion metrics associated with each campaign are primarily reported to demonstrate the variability in alpaca campaigns performed on the Beacon against different targets using different immunization strategies. Notably, even the same immunization regimen in multiple alpacas may result in a substantial degree of alpaca-to-alpaca variability in the antibody response^21^, suggesting that a relatively large number of alpacas might be required to determine an optimal immunization regimen for any given target or target class. To establish a robust alpaca immunization strategy, a carefully designed experiment needs to be performed using larger cohorts of alpacas for multiple immunogens.

## ACKNOWLEDGEMENTS

We thank Andrew Doucette, Kelly Rothenberger, Kirsten Grant, and Brooke McIsaac for performing titer tests on alpaca serum samples. We also acknowledge Jing Ouyang for helping with Beacon screen execution. We further thank Justin Stolte, Brendan Greamo, and Kumar Dasuri for performing transfections, harvests, and purifications of recombinant VHH-Fc clones, respectively. We also acknowledge Zsuzsanna Sükösd Etches, Max Gubert-Olivé, Kris Modig, Owen Bodley, and Jannick Bendtsen (affiliation: PipeBio ApS, Denmark) for their help with repertoire pipeline analysis and the associated figures. Finally, we thank Tracey Mullen, Aaron Sato, and Ryan Stafford (affiliation: Twist Bioscience Corporation) for their feedback on the manuscript.

## AUTHOR CONTRIBUTIONS

MBS, RK, and CAS designed the study, planned, and coordinated the PSMA immunizations and scheduling of harvests. MBS, AD, JG, and CAS designed experiments to develop and optimize alpaca cell enrichments, culturing conditions, and HCAb screening on the Beacon and performed Beacon screens. JB, KP, and CAS developed and optimized Sanger VHH sequencing, NGS repertoire sequencing, and data analysis. FG designed the humanization strategy and performed *in silico* bulk humanization of parental VHH-Fc clones. AB and VK designed, performed, and analyzed SPR and BLI experiments to characterize the parental VHH-Fc clones. JW coordinated the Target B immunizations and harvest scheduling. CAS, MBS, and VK designed and prepared the figures. MBS, CAS, AB, FG, JB, and JG wrote the manuscript. All authors reviewed the final version.

## FIGURE AND TABLE LEGENDS

Supplementary Table 1. PSMA binding kinetics for the VHH-mFc antibodies as measured by surface plasmon resonance.

