## Supplementary Table 1 for "Alpaca Single B Cell Interrogation and Heavy-Chain-Only Antibody Discovery on an Optofluidic Platform"

**Supplementary Table 1.** PSMA binding kinetics for the VHH-mFc antibodies as measured by surface plasmon resonance.

| Antibody ID | $k_a$ ($M^{-1}s^{-1}$) | $k_d$ ($s^{-1}$) | $K_D$ (M) | $t_{½}$ (min) |
|---|---|---|---|---|
| **D11D77826-2258** | 2.25E+04 | 9.14E-05 | 4.06E-09 | 126 |
| **D9D88792-8800** | 2.84E+04 | 9.25E-05 | 3.26E-09 | 125 |
| **D11D77826-1763** | 5.28E+03 | 3.32E-05 | 6.29E-09 | 348 |
| **D12D77826-11122** | 4.69E+03 | 2.59E-05 | 5.52E-09 | 446 |
| **D8D85764-3574** | 3.63E+03 | 1.33E-05 | 3.66E-09 | 868 |
| **D9D88792-5767** | 2.93E+03 | 3.40E-05 | 1.16E-08 | 340 |
| **D11D77826-3826** | 3.86E+04 | 1.69E-04 | 4.38E-09 | 68 |
| **D9D88792-8769** | 1.53E+04 | 1.93E-05 | 1.26E-09 | 598 |
| **D9D88792-13450** | 2.98E+04 | 6.29E-05 | 2.11E-09 | 184 |
| **D9D88792-14345** | 4.72E+03 | 1.86E-05 | 3.94E-09 | 621 |
| **D6D88695-15168** | 3.20E+03 | 1.11E-05 | 3.47E-09 | 1041 |
| **D9D88792-8908** | 1.23E+04 | 3.37E-05 | 2.74E-09 | 343 |
| **D11D77826-294** | 1.60E+04 | 6.54E-05 | 4.09E-09 | 177 |
